# Semantic overlap protects from interference by restoring encoding mechanisms

**DOI:** 10.1101/2022.02.07.479444

**Authors:** Isabelle L. Moore, Nicole M. Long

## Abstract

Overlap between experiences can have both facilitative and detrimental effects for memory. Our aim was to establish whether overlap along one dimension (e.g. contextual, semantic, temporal) can counteract overlap-driven interference along another dimension. We hypothesized that semantic overlap facilitates episodic memory formation by modulating encoding mechanisms. We recorded scalp electroencephalography (EEG) while human participants performed a free recall task. Half of the items from late in each study list semantically overlapped with an item presented earlier in the list. We find that semantic overlap selectively improves memory and influences the neural signals engaged during the study of late list items. Relative to other recalled items, late list items that are later recalled consecutively with semantically overlapping items elicited increased high frequency activity and decreased low frequency activity, a hallmark of successful encoding. Our findings demonstrate that semantic overlap can protect from interference due to temporal overlap by modulating encoding mechanisms.

**Statement of Relevanc:** Experiences can overlap along different dimensions, including contextual, semantic, and temporal. We tested the hypothesis that semantic overlap – shared meaning between experiences – may protect from interference due to temporal overlap, when experiences occur close together in time. Although evidence suggests that attention and/or encoding resources diminish across a series of study items presented in close temporal proximity, we find that semantic overlap can enable recovery of these encoding resources. Specifically, items that would typically be forgotten due to interference are better remembered and recruit distinct neural mechanisms when they share meaning with an earlier study item. These findings indicate that encoding mechanisms can be modulated by the degree of semantic overlap between two experiences. More broadly, our results suggest that experiences do not exist in isolation, rather that a prior experience can directly influence the neural mechanisms recruited to process a current experience.

## Introduction

In many situations, overlap between experiences produces interference (McGeoch, 1942; Anderson, 2003). Dining at two different Italian restaurants can impair memory for which dish was eaten at which restaurant. However, overlap can also facilitate inference judgments (Schlichting, Zeithamova, & Preston, 2014) and memory performance when freely recalling lists of words (Howard & Kahana, 2002). As experiences can overlap along many dimensions including contextual (Polyn, Norman, & Kahana, 2009), semantic (Howard & Kahana, 2002), and temporal (Smith, Moore, & Long, preprint), overlap may have different consequences depending on which dimension(s) is/are overlapping. The aim of the current study was to investigate whether overlap in one dimension can counteract overlap-driven interference in another dimension.

When studying a list of items, individuals can attend to either episodic features – when and where an item occurs – or semantic features – the meaning associated with an item. In free recall, reliance on episodic features is demonstrated through temporal clustering, the tendency to consecutively recall words that occupy neighboring positions on study lists (Kahana, 1996), and reliance on semantic features is demonstrated through semantic clustering, the tendency to consecutively recall words that share meaning (Bousfield, 1953). The general tendency to cluster responses during recall suggests that overlapping episodic or semantic features at study can facilitate subsequent memory, as clustering is typically positively correlated with overall recall (Sederberg, Miller, Howard, & Kahana, 2010; Healey, Crutchley, & Kahana, 2014). Likewise, we have previously found that the neural mechanisms of successful encoding – increased high frequency spectral activity (signals above 28 Hz) and decreased low frequency spectral activity (signals below 28 Hz) in left prefrontal cortex and hippocampus (Long, Burke, & Kahana, 2014) – are also recruited during the study of items that will subsequently be temporally or semantically clustered (Long & Kahana, 2015, 2017).

Despite the apparent memory benefits described above, overlapping features can also lead to decreased memory performance in free recall. The primacy effect in delayed recall – better memory for early relative to late list items (Murdock, 1962) – may be due to an attentional decline (Azizan & Polich, 2007) and/or depletion of encoding resources (Lohnas, Davachi, & Kahana, 2020, ‘neural fatigue’) across items. In a typical study list, words do not overlap semantically, meaning that the only available features are episodic features. Potentially, attention to episodic features may decline as the list proceeds, leading to worse memory. Similarly, overlapping semantic features across study lists can produce a build up of proactive interference, whereby performance declines across lists (Underwood, 1957; Watkins & Watkins, 1975; Postman & Keppel, 1977; Szpunar, McDermott, & Roediger, 2008). In both of these instances, study items overlap in either the temporal or semantic dimension, but not both. If attention to a given feature dimension declines during study, the introduction of a new overlapping feature could potentially recover encoding processes and improve memory performance.

Our hypothesis is that overlap along one dimension can protect from interference due to overlap in another dimension by modulating encoding mechanisms. Here, we specifically test whether semantic overlap can facilitate episodic memory formation. We conducted a human scalp electroencephalography (EEG) study in which participants performed a free recall task. We manipulated the degree to which ‘pairs’ of individually presented words were semantically overlapping while controlling the serial position of each word. Participants studied pairs of strongly semantically associated words (e.g. ‘dog’ and ‘cat’) and pairs of weakly semantically associated words (e.g. ‘shore’ and ‘road’). The second word in each of these pairs (cat, road) appeared in a later serial position compared to the first word in each pair (dog, shore). Due to decreasing attention to episodic features, memory for second words should be worse than for first words, but we hypothesize that this memory decline may be ameliorated if the second word overlaps semantically with the first word. We separately measured behavioral performance and neural signals for the first and second words in each pair. To the extent that the encoding of second words is influenced by semantic overlap with a first word, we expect to find differential memory performance and neural activity patterns for words that overlap semantically, compared to those that do not.

## Materials and Methods

### Participants

Forty (22 female; mean age 20.63 years) native English speakers from the University of Virginia community participated. We selected a sample size of N = 40 based on other scalp EEG studies conducted in our lab (Long & Kuhl, 2019; Smith et al., preprint). All participants had normal or corrected-to-normal vision. Informed consent was obtained in accordance with the University of Virginia Institutional Review Board for Social and Behavioral Research and participants were compensated for their participation. Two participants were excluded from the final dataset: one whose EEG recording was not started until the third run, and one whose verbal responses were not recorded. Thus, data are reported for the remaining 38 participants. The raw, de-identified data and the associated experimental and analysis codes used in this study will be made available via the Long Term Memory laboratory website upon publication.

### Experimental Design and Statistical Analysis

#### Free Recall Task

Stimuli consisted of 1602 words, drawn from the Toronto Noun Pool (Friendly, Franklin, Hoffman, & Rubin, 1982). From this set, 192 words were randomly selected for each participant. Words were presented in lists of 16 words across a total of 12 runs.

#### Study phase

During each trial, participants viewed a single word presented for 2000 ms followed by a 1000 ms inter-stimulus interval (ISI; Fig. 1). Participants were instructed to study the presented word in anticipation of a later memory test; participants did not make any behavioral responses. Each list was comprised of 16 words split into two parts (“first associates” and “second associates,” respectively) separated by a brief 2000 ms delay and a get ready screen. The critical manipulation was the strength of semantic association between first and second associates. Semantic association strength was determined using Word Association Space values (WAS; Nelson, Zhang, & McKinney, 2001); ‘strong’ semantic associates had a WAS value of 0.4 or greater and ‘weak’ semantic associates had a WAS value less than 0.4 (Long & Kahana, 2017). Each first associate was “paired” with a second associate and separated by seven intervening items (a lag of eight). As an example, in Fig. 1, the dog-cat pair is comprised of strong semantic associates (WAS = 0.86); dog is the first associate and cat is the second associate. In comparison, the shore-road pair is comprised of weak semantic associates (WAS = 0.017). Both strong and weak semantic associates were weakly semantically associated to all other study words.

**Fig. 1.**
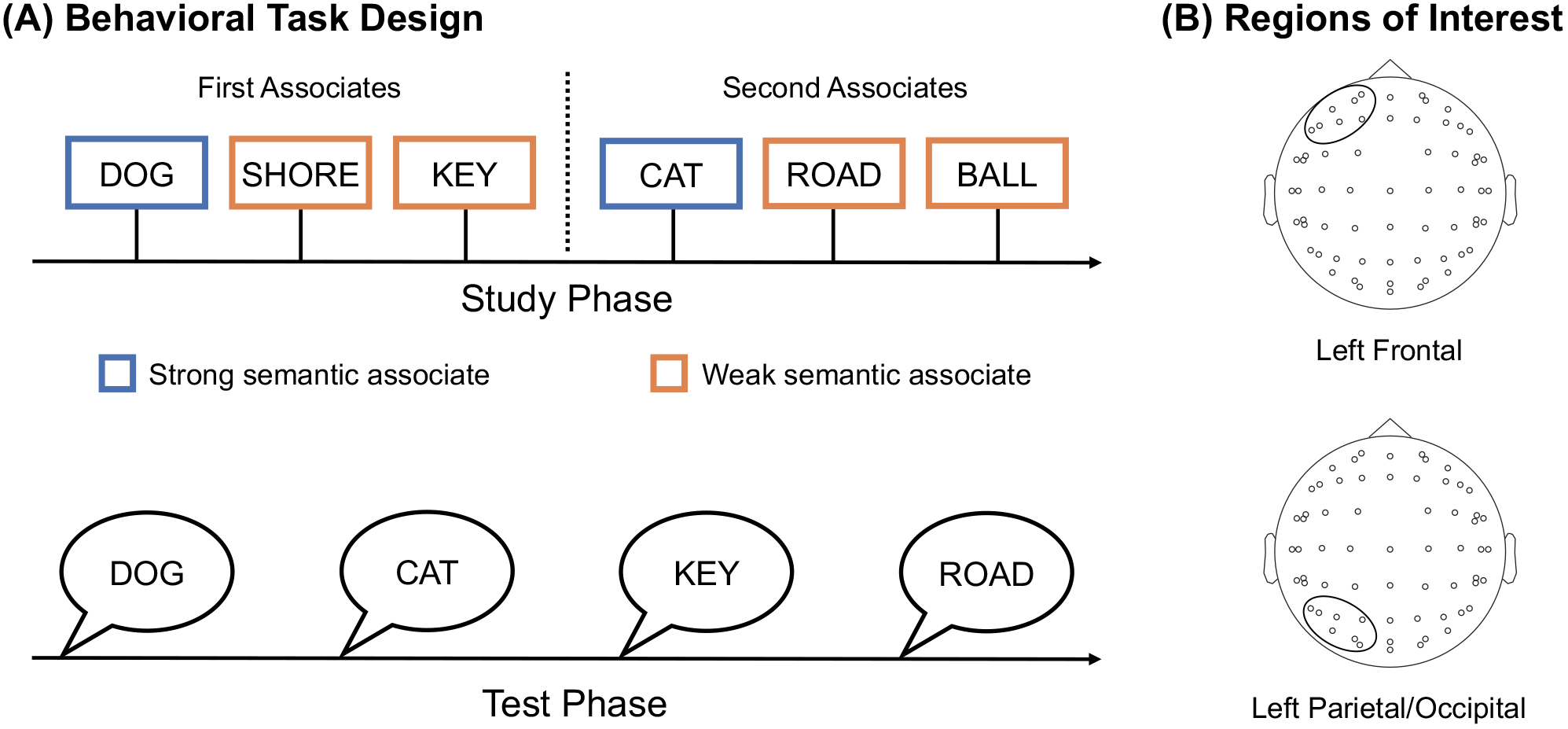
Experiment methods. **(A)** During the study phase, participants studied a series of words one at a time in anticipation of a later memory test. Each word list was split into “first associates” (e.g. “dog,” “shore,” “key”) and “second associates” (e.g. “cat,” “road,” “ball”) which were paired with one another. Half of the associate pairs were strongly semantically associated (e.g. “dog” and “cat”) and the other half were weakly semantically associated (e.g. “shore” and “road”). Strong semantic associates are shown here in blue and weak semantic associates are shown in orange for demonstration purposes only; participants were not given any indication of semantic association strength. During the test phase, participants verbally recalled any words that they could remember from the immediately preceding study phase, in any order. **(B)** Regions of interest (ROIs): We analyzed two ROIs, left frontal and left parietal/occipital.

#### Math distractor phase

On each trial, participants saw a three digit math problem with a solution (of the form, “X + Y - Z = A”). Participants had 4 seconds to verify whether the solution shown was correct. Each math problem was followed by a minimum 1000 ms ISI. If a response was made under 4 seconds then the ISI was 1000 ms plus the remaining time. Participants saw a total of four math problems, randomly generated, such that the distractor interval was always 20 seconds in duration.

#### Free recall phase

Following the math distractor, an auditory beep cued the participant to verbally recall any words that they could remember from the immediately preceding study phase. Participants were given 45 seconds to recall as many words as possible in any order. Participants were encouraged to continue trying to recall throughout the interval.

### EEG data acquisition and preprocessing

EEG recordings were collected using a BrainAmp system (Brain Products, Inc.) and an ActiCap equipped with 64 Ag-AgCl active electrodes positioned according to the extended 10-20 system. All electrodes were digitized at a sampling rate of 1000 Hz and were referenced to electrode FCz. Offline, electrodes were later converted to an average reference. Impedances of all electrodes were kept below 50kΩ. Electrodes that demonstrated high impedance or poor contact with the scalp were excluded from the average reference. Bad electrodes were determined by voltage thresholding (see below).

Custom Python codes were used to process the EEG data. We applied a high pass filter at 0.1 Hz, followed by a notch filter at 60 Hz and harmonics of 60 Hz to each participant’s raw EEG data. We then performed three preprocessing steps (Nolan, Whelan, & Reilly, 2010) to identify and correct electrodes with severe artifacts separately for each participant. First, we calculated the mean correlation between each electrode and all other electrodes as electrodes should be moderately correlated with other electrodes due to volume conduction. We *z*-scored these means across electrodes and rejected electrodes with *z*-scores less than −3. Second, we calculated the variance for each electrode as electrodes with very high or low variance across a session are likely dominated by noise or have poor contact with the scalp. We then *z*-scored variance across electrodes and rejected electrodes with a *|z| ≥* 3. Finally, we expect many electrical signals to be autocorrelated, but signals generated by the brain versus noise likely have different forms of autocorrelation. Therefore, we calculated the Hurst exponent, a measure of long-range autocorrelation, for each electrode and rejected electrodes with a *|z| ≥* 3. Rejected electrodes were excluded from the average re-reference. We found the average voltage across all of the remaining electrodes for each time sample and re-referenced the data by subtracting the average voltage from the filtered EEG data. We used wavelet-enhanced independent component analysis (Castellanos & Makarov, 2006) to remove artifacts from eyeblinks and saccades.

### EEG data analysis

In order to perform spectral decomposition, we applied a family of Morlet wavelet transforms (wave number = 6) to all electrode EEG signals across 46 logarithmically-spaced frequencies (2-100 Hz; Long & Kahana, 2015). After log-transforming the power, we downsampled the data by taking a moving average across 100 ms time intervals from −4000 to 4000 ms relative to stimulus onset and sliding the window every 25 ms, resulting in 317 time intervals (80 nonoverlapping). Power values were then *z*-transformed by subtracting the mean and dividing by the standard deviation power. Mean and standard deviation power were calculated across all trials and across time points for each frequency. We divided the *z*-transformed power (zPower) into six frequency bands: low theta (3-4 Hz), high theta (6-8 Hz), alpha (10-14 Hz), beta (16-26 Hz), low gamma (28-42 Hz), and high gamma (44-100 Hz; Long & Kahana, 2017).

### Regions of Interest

We selected two regions of interest (ROIs), left frontal (Fp1, F3, F7, AF7, AF3, F1, F5) and left parietal/occipital (P3, P7, O1, P1, P5, PO7, PO3), based on our prior work (Long & Kahana, 2017). We focused on the left hemisphere given evidence that subsequent memory effects are typically left lateralized (Kim, 2011; Burke et al., 2014).

### Behavioral data analysis

We assessed study items based on associate (first or second) and semantic association strength (strong or weak). Strong semantic associates could be subsequently recalled and semantically clustered (SClust), whereby the study item was recalled preceding or following its semantic associate. By definition, weak semantic associates could not be semantically clustered. Any study item could be subsequently recalled and not clustered (NClust), whereby the study item was recalled, but not consecutively with either a study neighbor or a semantic associate. We assessed the tendency of participants to semantically cluster their responses by performing a semantic conditional response probability (sCRP) analysis (Howard & Kahana, 2002), in which we calculated the probability of recalling an item as a function of having just recalled an item with a given level of semantic association strength (WAS value) to the current item. We grouped words into 6 semantic association strength bins based on WAS values: below 0, 0 to 0.2, 0.2 to 0.4, 0.4 to 0.6, 0.6 to 0.8, and 0.8 to 1.0, where ‘below 0’ is the lowest semantic association strength bin and ‘0.8 to 1.0’ is the highest semantic association strength bin. We also reduced the sCRP to a single semantic clustering score by finding the difference between the average sCRP value for the low semantic association strength bins (below 0 - 0.4) and the high semantic association strength bins (0.4 - 1.0) for each participant (Long & Kahana, 2017).

### Univariate data analysis

We performed two univariate contrasts. First, we compared spectral signals during the study of strong second associates that were subsequently semantically clustered (SClust_*s*_) and weak second associates that were subsequently recalled but not clustered (NClust_*w*_). Second, we compared the semantic subsequent clustering effect (SCE_*s*_) between first and second strong associates. The SCE_*s*_ is the difference in zPower between subsequently recalled associates that are vs. are not semantically clustered. For each contrast, participant, electrode, and frequency, we calculated zPower in each of the two conditions, averaged over the 2000 ms stimulus interval for each ROI.

### Statistical analyses

We used paired-sample *t*-tests and a repeated measures ANOVA (rmANOVA) to assess the effects of semantic association strength (strong, weak) and associate (first, second) on probability of recall. We used an rmANOVA to assess the effects of semantic association strength (strong, weak) and serial position (1-16) on probability of recall. We used paired-sample *t*-tests to compare the conditional response probability for strong and weak semantic bins. We used a Pearson correlation to measure the relationship between semantic clustering score and probability of recall for each associate type (strong, weak *×* first, second). We used rmANOVAs to assess the effects of subsequent clustering condition (SClust_*s*_, NClust_*w*_) and frequency on zPower. We used rmANOVAs to assess the effects of associate and frequency on the SCE_*s*_.

## Results

### Influence of semantic associations on free recall

According to our hypothesis, semantic overlap may protect late list items from interference. Thus, our first goal was to test whether first and second associates are differentially remembered by virtue of their semantic association strength. We ran a two-way, rmANOVA to evaluate the effects of semantic association strength (strong, weak) and associate (first, second) on probability of recall (Fig. 2A). We found a main effect of semantic association strength (*F*_1,37_ = 29.16, *p* < 0.001, *η_p_*^2^ = 0.44) driven by greater probability of recall for strong than weak semantic associates. We found a main effect of associate (*F*_1,37_ = 8.46, *p* = 0.006, *η_p_*^2^= 0.19) driven by greater probability of recall for the first associate compared to the second associate. We found a significant interaction between semantic association strength and associate (*F*_1,37_ = 8.48, *p* = 0.006, *η_p_*^2^ = 0.19). The difference in probability of recall between strong and weak semantic associates was greater for second (*M* = 0.10, *SD* = 0.10) compared to first (*M* = 0.04, *SD* = 0.09) associates (*t*_37_ = 2.91, *p* = 0.006, *d* = 0.57). These results demonstrate that semantic association strength differentially affects probability of recall of early vs. late list items. Late list items are typically remembered less well than early list items in delayed free recall, potentially due to a build up of interference. Our behavioral findings support our hypothesis that late list items may be protected from this interference by virtue of semantic overlap with an early list item.

**Fig. 2.**
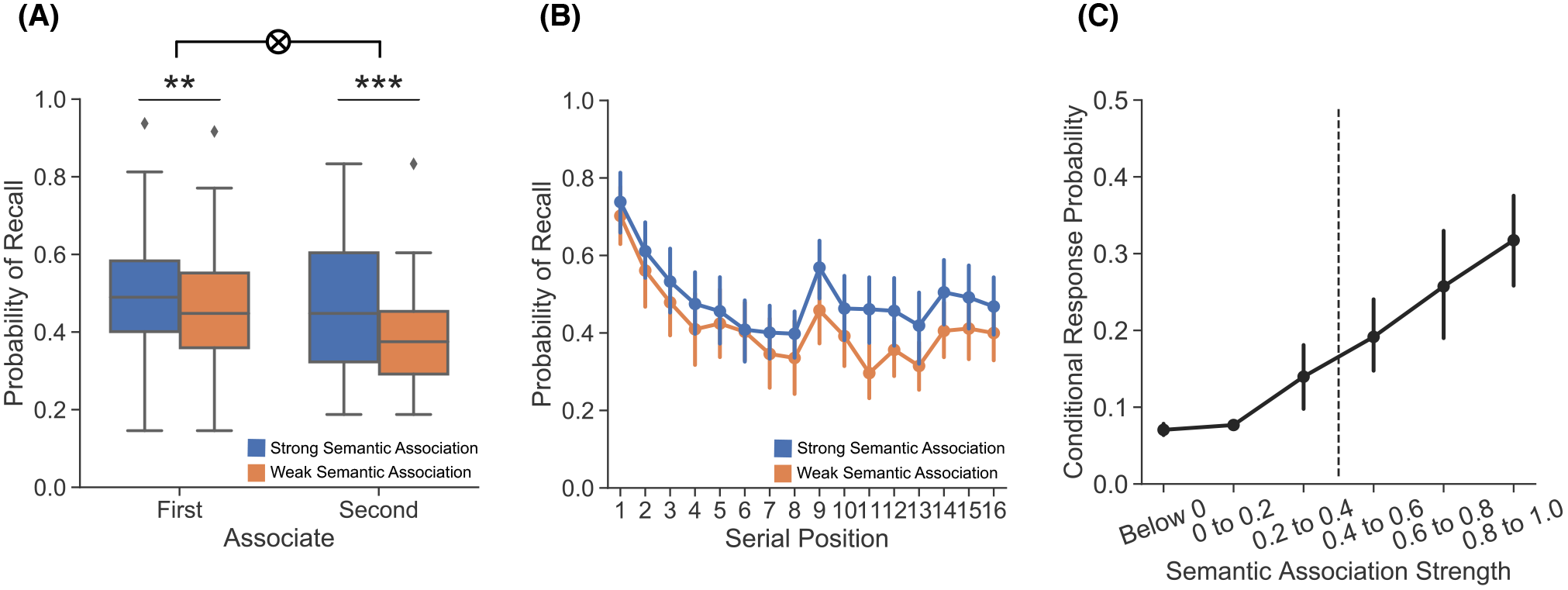
Recall performance and memory organization. **(A)** Probability of recall was greater for strong (blue) compared to weak (orange) semantic associates and was greater for first compared to second associates. There was a significant interaction between associate and semantic association strength (*p* = 0.006). **(B)** Probability of recall was greater for strong than weak semantic associates across serial positions. **(C)** Participants were more likely to make transitions between strong compared to weak semantic associates. The vertical, dashed line denotes the boundary between weak semantic bins (below 0, 0.0-0.2, 0.2-0.4) and strong semantic bins (0.4-0.6, 0.6-0.8, 0.8-1.0). Error bars denote 95% confidence intervals. *** p < 0.001, ** p < 0.01.

Although probability of recall is greater for strong relative to weak second associates, the previous analysis does not indicate whether this increase is consistent across *all* strong second associates. We expect all semantically overlapping stimuli to be protected from interference; however, semantic overlap may selectively increase the salience of initially presented strong second associates, an effect that could diminish as participants encounter additional strong second associates. To adjudicate between these possibilities, we measured probability of recall as a function of semantic association strength and serial position. We ran a 2 × 16 rmANOVA to evaluate the effects of semantic association strength (strong, weak) and serial position (1-16) on probability of recall (Fig. 2B). We found a main effect of semantic association strength (*F*_1,37_ = 27.76, *p* < 0.001, *η_p_*^2^ = 0.43) driven by greater probability of recall for strong than weak semantic associates. We found a main effect of serial position (*F*_15,555_ = 13.67, *p* < 0.001, *η_p_*^2^ = 0.27). We did not find an interaction between semantic association strength and serial position (*F*_15,555_ = 0.647, *p* = 0.836, *η_p_*^2^ = 0.02). Bayes factor analysis revealed that a model without the two-way interaction term is preferred to a model with the two-way interaction by a factor of 3117.32. The lack of an interaction between association strength and serial position suggests that semantic overlap protects all overlapping late list items from interference, regardless of serial position.

Our next goal was to directly link the recall improvement for strong second associates to semantic processing of those items. If semantic processing yields greater probability of recall specifically for strong second associates, then (1) participants should show a tendency to semantically cluster their recalls and (2) the degree to which a participant clusters their recalls should be positively correlated with probability of recall selectively for strong second associates. We measured semantic clustering by performing a semantic conditional response probability (sCRP) analysis (Howard & Kahana, 2002). Briefly, the sCRP analysis reveals the overall tendency to consecutively recall two items on the basis of their semantic association strength. We defined three strong and three weak semantic association strength bins based on WAS values (see Methods). Participants are more likely to make transitions between strong (*M* = 0.26, *SD* = 0.19) compared to weak semantic associates (*M* = 0.10, *SD* = 0.08; *t*_37_ = 9.15, *p* < 0.001; Fig. 2C). We reduced the sCRP to a single semantic clustering score (average transition probability for strong bins - average transition probability for weak bins; Long & Kahana, 2017), and found that there is a positive across-participant correlation between semantic clustering and memory for strong second associates (*r*_37_ = 0.366, *p* = 0.024), but no such association for memory for the other associates (strong first, *r*_37_ = 0.230, *p* = 0.164; weak first, *r*_37_ = 0.074, *p* = 0.660; weak second, *r*_37_ = 0.228, *p* = 0.168). These results indicate that higher levels of semantic clustering are associated with better recall for strong second associates.

### Dissociable neural substrates for semantically clustered second associates

We hypothesized that semantic overlap alters the encoding mechanisms recruited during the study of late list items. To test this hypothesis, we compared spectral power during the study of strong second associates that were subsequently semantically clustered (SClust_*s*_) and weak second associates that were subsequently recalled but not clustered (NClust_*w*_). We selected this contrast as it holds constant the serial position of the items and the overall memory for the items c– both conditions include only second associates that are subsequently recalled – while varying the potential influence of semantic overlap and consequent semantic processing on those items. To the extent that semantic overlap influences encoding mechanisms for late list items, the spectral signals during SClust_*s*_ vs. NClust_*w*_ items will differ. We ran a 2 *×* 6 rmANOVA to evaluate the effects of condition (SClust_*s*_, NClust_*w*_) and frequency on zPower separately for our two ROIs (Fig. 3A). Over left frontal (LF), we found a main effect of frequency (*F*_5,185_ = 3.121, *p* = 0.010, *η_p_*^2^ = 0.08), no main effect of condition (*F*_1,37_ = 0.334, *p* = 0.567, *η_p_*^2^ = 0.01) and a significant interaction between condition and frequency (*F*_5,185_ = 4.193, *p* = 0.001, *η_p_*^2^ = 0.10).

**Fig. 3.**
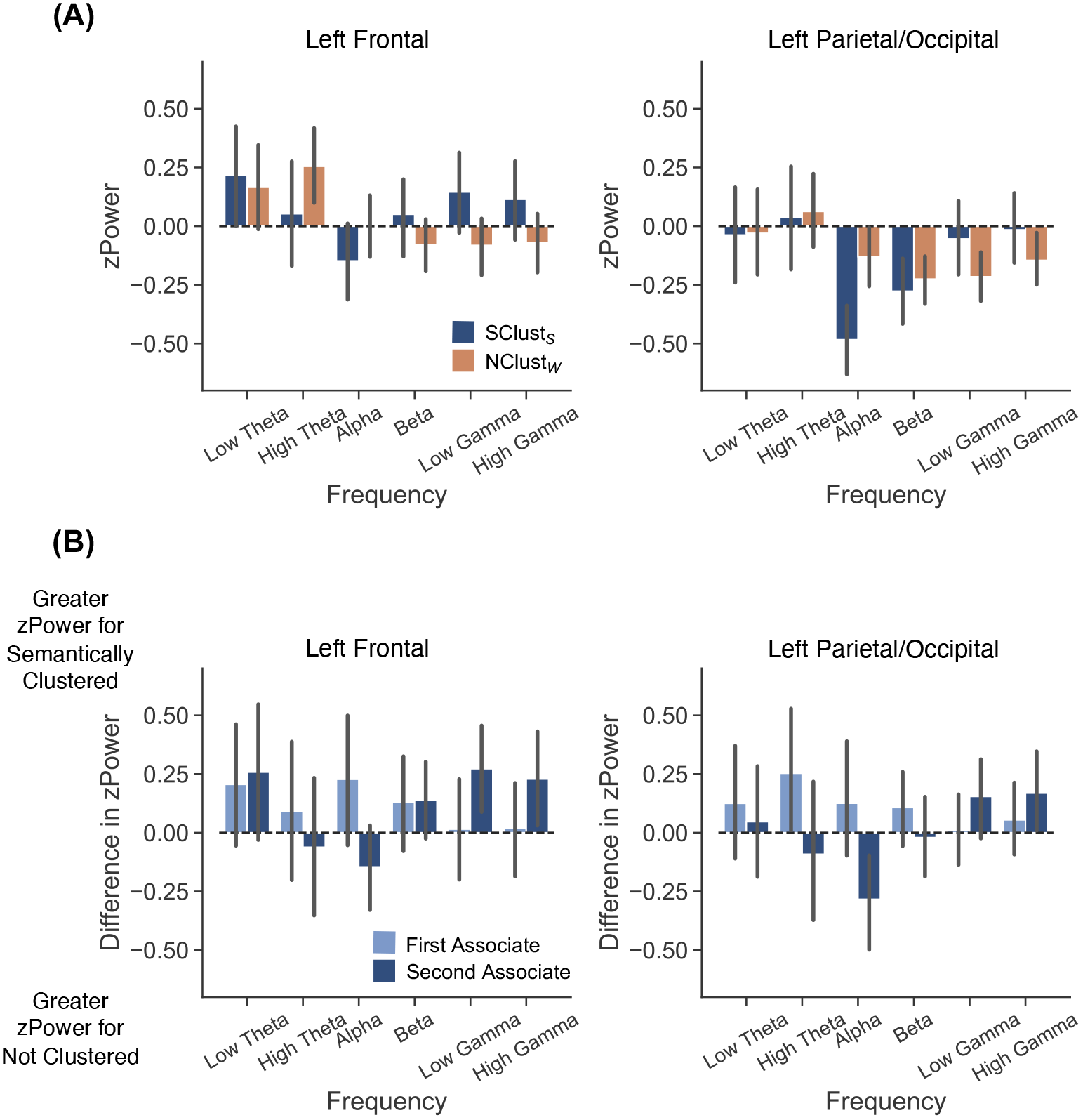
Differential univariate effects for semantically clustered second associates. zPower is shown for six frequency bands (low theta: 3-4 Hz, high theta: 6-8 Hz, alpha: 10-14 Hz, beta: 16-26 Hz, low gamma: 28-42 Hz, high gamma: 44-100 Hz) across two regions of interest, left frontal and left parietal/occipital. zPower is averaged across the stimulus interval (2000 ms). Error bars denote 95% confidence intervals. **(A)** The top panel shows zPower for second associates that were strongly semantically associated and subsequently semantically clustered (blue, SClust_*s*_) and second associates that were weakly semantically associated and subsequently recalled but not clustered (orange, NClust_*w*_). **(B)** The bottom panel shows the difference in zPower between strong items that were subsequently semantically clustered and strong items that were subsequently recalled, but not clustered, for first (light blue) and second associates (dark blue). Positive values indicate greater zPower for items later semantically clustered, negative values indicate greater zPower for items later recalled, but not clustered.

Over left parietal/occipital (LP), we found a main effect of frequency (*F*_5,185_ = 5.154, *p* < 0.001, *η_p_*^2^ = 0.12), no main effect of condition (*F*_1,37_ = 0.35, *p* = 0.558, *η_p_*^2^ = 0.01) and a significant interaction between condition and frequency (*F*_5,185_ = 3.742, *p* = 0.003, *η_p_*^2^ = 0.09). Relative to NClust_*w*_ items, SClust_*s*_ items showed increased high frequency activity (HFA_*i*_; signals above 28Hz) and decreased low frequency activity (LFA_*d*_; signals below 28Hz). These results suggest that late list items are processed differently based on their semantic overlap with early list items.

The difference in neural signals between strong vs. weak second associates may be the result of either semantic-specific processing or a more general binding mechanism that links items to their spatiotemporal context (Howard & Kahana, 2002; Sederberg, Howard, & Kahana, 2008; Polyn et al., 2009; Lohnas & Kahana, 2014). During study, the HFA_*i*_/LFA_*d*_ pattern predicts both subsequent memory (Sederberg et al., 2006; Burke et al., 2013; Long et al., 2014) and subsequent clustering (Long & Kahana, 2015, 2017) and is thought to reflect item-context binding. Hence, the dissociation that we observe between second associates may be driven by increased item-context binding during strong compared to weak associates, rather than selective semantic processing of the strong associates. To test these alternatives, we directly compared the subsequent semantic clustering effect (SCE_*s*_) across first and second strong associates. The SCE_*s*_ is the difference in zPower between strong associates that are subsequently semantically clustered (SClust_*s*_) and strong associates that are subsequently recalled but not clustered (NClust_*s*_). If the SCE_*s*_ differs between first and second associates, this would support our hypothesis that semantic overlap differentially impacts late list items. If the SCE_*s*_ is the same across first and second associates, this would support the alternative interpretation that a general item-context binding mechanism underlies encoding of both early and late list items that are subsequently clustered. We ran a 2 *×* 2 *×* 6 rmANOVA to evaluate the effects of associate (first, second), condition (SClust_*s*_, NClust_*s*_), and frequency on zPower (Fig. 3B). A two-way interaction between condition and frequency, with no three-way interaction between condition, frequency, and associate, would indicate that the SCE_*s*_ is the same for both first and second associates. A three way interaction would indicate that the SCE_*s*_ differs across first and second associates. We report the results of this ANOVA in Table 1 and highlight the key findings below.

**Table 1.** Analysis of variance (ANOVA) results for associate, frequency, and condition on zPower.

| Effect | Left Frontal |  |  |  | Left Parietal/Occipital |  |  |  |
| --- | --- | --- | --- | --- | --- | --- | --- | --- |
| | df | *F* | *p* | $\eta_p^2$ | df | *F* | *p* | $\eta_p^2$ |
| Main effect of frequency | (5,185) | 1.45 | 0.208 | 0.04 | (5,185) | 14.62 | < **0.001** | 0.28 |
| Main effect of associate | (1,37) | 0.17 | 0.686 | 0.00 | (1,37) | 7.62 | **0.009** | 0.17 |
| Main effect of condition | (1,37) | 6.20 | **0.017** | 0.14 | (1,37) | 2.52 | 0.121 | 0.06 |
| Interaction of condition × associate | (1,37) | 0.00 | 0.995 | 0.00 | (1,37) | 2.55 | 0.119 | 0.06 |
| Interaction of condition × frequency | (5,185) | 0.951 | 0.449 | 0.03 | (5,185) | 0.79 | 0.558 | 0.02 |
| Interaction of associate × frequency | (5,185) | 1.71 | 0.134 | 0.04 | (5,185) | 5.66 | < **0.001** | 0.13 |
| Interaction of condition × associate × frequency | (5,185) | 3.15 | **0.009** | 0.08 | (5,185) | 2.33 | **0.044** | 0.06 |

Over both LF and LP, we find a three-way interaction between associate, frequency and condition (LF: *F*_5,185_ = 3.15, *p* = 0.009, *η_p_*^2^ = 0.08; LP: *F*_5,185_ = 2.33, *p* = 0.044, *η_p_*^2^ = 0.06) and no interaction between condition and frequency (LF: *F*_5,185_ = 0.95, *p* = 0.449, *η_p_*^2^ = 0.03; LP: *F*_5,185_ = 0.79, *p* = 0.558, *η_p_*^2^ = 0.02). These results demonstrate that the SCE_*s*_ varies across first and second associates, providing support for our hypothesis that semantic overlap differentially impacts late list items.

The SCE_*s*_ dissociation across first and second associates indicates that semantic overlap impacts the neural signals during the study of late list items. However, as our hypothesis is that semantic overlap specifically alters encoding mechanisms, we should *selectively* observe the SCE_*s*_ for late list items and the SCE_*s*_ for those items should exhibit the HFA_*i*_/LFA_*d*_ pattern. Within each ROI, we ran a 2 *×* 6 rmANOVA to evaluate the effects of condition (SClust_*s*_, NClust_*s*_) and frequency on zPower. For first associates, we found no main effect of condition (LF: *F*_1,37_ = 1.896, *p* = 0.177, *η_p_*^2^ = 0.05; LP: *F*_1,37_ = 3.835, *p* = 0.058, *η_p_*^2^ = 0.09), a main effect of frequency in LP (LF: *F*_5,185_ = 1.896, *p* = 0.177, *η_p_*^2^ = 0.05; LP: *F*_5,185_ = 11.27, *p* < 0.001, *η_p_*^2^ = 0.23), and no interaction between condition and frequency (LF: *F*_5,185_ = 0.758, *p* = 0.581, *η_p_*^2^ = 0.02; LP: *F*_5,185_ = 0.627, *p* = 0.679, *η_p_*^2^ = 0.02). Bayes factor analysis revealed that a model without the two-way interaction term is preferred to a model with the two-way interaction by a factor of 55.08 (LF) and 57.19 (LP). For second associates, we found no main effect of condition (LF: *F*_1,37_ = 3.11, *p* = 0.086, *η_p_*^2^ = 0.08; LP: *F*_1,37_ = 0.01, *p* = 0.919, *η_p_*^2^ = 0.00), a main effect of frequency in LP (LF: *F*_5,185_ = 1.342, *p* = 0.249, *η_p_*^2^ = 0.04; LP: *F*_5,185_ = 8.416, *p* < 0.001, *η_p_*^2^ = 0.19), and a significant interaction between condition and frequency (LF: *F*_5,185_ = 2.916, *p* = 0.015, *η_p_*^2^ = 0.07; LP: *F*_5,185_ = 2.314, *p* = 0.046, *η_p_*^2^ = 0.06). That we find an SCE_*s*_ for second associates, but failed to find an SCE_*s*_ for first associates, provides support for our hypothesis that semantic overlap modulates encoding mechanisms. Importantly, we find that this selective SCE_*s*_ is characterized by the HFA_*i*_/LFA_*d*_ pattern – a marker of successful encoding – suggesting that semantic overlap can enhance encoding for items that may otherwise be poorly encoded due to their position in the study list.

## Discussion

The goal of the current study was to investigate the extent to which multiple forms of overlap interact to protect from interference. We recorded scalp EEG while participants performed a free recall task in which each study list was comprised of words that did and did not overlap semantically. We report four key findings. First, we find that semantic overlap with a prior list item improves memory for items presented later in the study list. Second, we find that participants’ tendency to semantically cluster is associated with an increased likelihood of specifically recalling semantically overlapping late list items. Third, we find differential spectral signals during late list item encoding; subsequently semantically clustered items are distinct from non-overlapping items that are subsequently recalled but not clustered. Finally, we find selective high frequency activity increases and low frequency activity decreases – a hallmark of successful encoding – for late list items that are semantically clustered, compared to early list items that are semantically clustered. Taken together, these results suggest that semantic overlap can modulate encoding mechanisms to protect from interference due to temporal overlap.

Late list items are better remembered if they semantically overlap with an early list item. In delayed free recall tasks in which no words are semantically associated, recall performance decreases over serial position (Murdock, 1962). This decrease in probability of recall may arise from decreases in attention (Sederberg et al., 2006; Azizan & Polich, 2007), and/or a decline in encoding resources (Lohnas et al., 2020; Popov & Reder, 2020; Mizrak & Oberauer, 2021). Consistent with previous research, we find a decrease in probability of recall for late list items that did not semantically overlap with a prior list item. However, we find that semantic overlap counteracts this decrease in probability of recall, such that recall is increased for semantically overlapping late list items regardless of serial position. Our interpretation is that as a list progresses, attention to episodic features declines, but that semantic overlap reorients attention to semantic features. Potentially, as more events become linked to the same spatiotemporal context, spatiotemporal features may become less salient or diagnostic (Nairne, 2002), leading to a decrement in memory formation. It is possible that semantically overlapping items are particularly surprising or distinctive, and that this accounts for the improved recall for these items.

However, according to this interpretation, items should be less surprising both as the study list and the experiment as a whole progresses, yet we find a benefit for semantically overlapping items across all serial positions. Thus our results support the interpretation that semantic overlap may specifically direct attention to the semantic features of an event.

In line with the interpretation that participants selectively attend to semantic features, participants semantically clustered their responses, and in doing so, were specifically more likely to recall late list items which overlapped semantically with early list items. Although temporal clustering has repeatedly been positively associated with probability of recall (Sederberg et al., 2010; Healey et al., 2014), the link between semantic clustering and recall performance is less clear, given that study lists are typically comprised of words that are not semantically associated. Indeed, attending to semantic features in the absence of semantically overlapping items can be detrimental for later memory (Long & Kahana, 2017), indicating that *de facto* reliance on semantic features is not adaptive. Our results suggest that attending to semantic features can be selectively beneficial when items that might otherwise be forgotten due to their serial position semantically overlap with a prior list item.

Semantic overlap selectively alters the encoding mechanisms recruited during the study of late list items. Specifically, we find that semantically clustered late list items are characterized by an increase in high frequency activity (HFA_*i*_; signals above 28Hz) and a decrease in low frequency activity (LFA_*d*_; signals below 28Hz), both when compared to non-overlapping late list items that are recalled but not clustered, and when compared to semantically clustered early list items. As the HFA_*i*_/LFA_*d*_ pattern predicts both subsequent memory (Long et al., 2014) and subsequent clustering (Long & Kahana, 2015, 2017), it may reflect the degree to which items are bound to their spatiotemporal context (Polyn et al., 2009). However, as the HFA_*i*_/LFA_*d*_ pattern is prevalent outside of the domain of memory (Crone, Boatman, Gordon, & Hao, 2001; Bauer, Oostenvald, Peeters, & Fries, 2006; Lachaux et al., 2007; Dalal et al., 2009), it may instead reflect task or attentional demands rather than memory *per se* (Long & Kuhl, 2019). That we observe HFA_*i*_/LFA_*d*_ selectively for overlapping late items is most consistent with the latter account. The dissociations that we currently observe cannot reflect a difference between subsequent remembering and subsequent forgetting as our contrasts only include items that were subsequently recalled. Furthermore, the dissociations cannot reflect a general item-context binding mechanism promoting subsequent semantic clustering as subsequently semantically clustered early list items did not show the HFA_*i*_/LFA_*d*_ pattern. Instead, the selectivity of the HFA_*i*_/LFA_*d*_ pattern is consistent with the interpretation that semantic overlap leads to a reorienting of attention toward semantic features and away from temporal features. Such a reorientation is not possible for non-overlapping late list items, or for overlapping early list items, as participants cannot know which early list items will be paired with an overlapping item later in the list. At its core, our interpretation is that semantic overlap changes *how* an item is encoded, specifically by increasing the attention directed to the semantic features of an experience.

Together, these results show that overlap along one dimension can protect from interference due to overlap in another dimension by modulating encoding mechanisms. An exciting direction for future research will be to investigate the extent to which these effects generalize to other dimensions beyond semantic overlap. Our findings demonstrate that study-phase mechanisms are not static, but instead item processing is influenced by prior experiences. More broadly, we contribute to a growing body of literature characterizing the facilitative and detrimental effects of overlapping experiences on cognition.

## Acknowledgments

We thank Yuju Hong for assistance with data collection. Nicole Long is an iTHRIV Scholar. The iTHRIV Scholars Program is supported in part by the National Center for Advancing Translational Sciences of the National Institutes of Health under Award Numbers UL1TR003015 and KL2TR003016.

